# Chromosomal genomes assemblies of twelve sheep breeds

**DOI:** 10.64898/2026.10.08.757562

**Authors:** Eché Camille, Birbes Clément, Marcuzzo Camille, Sarry Julien, Suin Amandine, Vernette Caroline, Kuchly Claire, Gaspin Christine, Milan Denis, Donnadieu Cécile, Iampietro Carole, Tosser-Klopp Gwenola, Klopp Christophe

## Abstract

The sheep research community benefits from a genome assembly that has been refined over the last 15 years. However, the rich diversity of sheep breeds suggests that a single individual genome is not enough to represent the species full genomic content. Third-generation sequencing technologies have recently made it possible to generate high quality assemblies that should enable to elucidate unique and/or complex phenotype determinisms depending on genetic structural variations. In the SeqOccin project, twelve breeds were selected to represent (mostly French) sheep genetic diversity: Basco-Béarnaise, Black Belly, Blanche du Massif Central, Corse, Ile de France, Lacaune, Manech Tête Noire, Noire du Velay, Ouessant, Rambouillet, Vendéen, Soay. High-molecular-weight DNA was extracted from blood samples and subjected to long read whole-genome sequencing. The reads were used to generate *de novo* assemblies and were aligned onto the public reference to call structural variants. PacBio HiFi and parental Illumina reads were used to assemble phased haplotypes of the Lacaune breed. For the other breeds, PacBio raw (CLR) or ONT reads were generated. Our work demonstrates the feasibility of assembling genomes with a reasonable amount of data and appropriate software packages, delivering one high-quality assembly and 11 standard sheep assemblies that pave the way for pangenome construction and structural variant functional studies.

## Introduction

Sheep (Ovis aries) were domesticated around 10,000 years ago in the Fertile Crescent in contemporary Iran (Dobney & Larson, 2006; Zeder, 2008), shortly after goats. Since then, they have spread worldwide, adapting to diverse environments and production systems. Through artificial and natural selection, sheep breeds have been specialised for the production of meat, milk, and wool (FAO, 2015) (http://www.fao.org/3/i5077e/i5077e.pdf), making them one of the most versatile and economically important livestock species. According to FAOSTAT (www.fao.org/faostat/en), the global sheep population is estimated to exceed 1.5 billion animals, with significant increases in Africa and Asia over the past decades.

In France, around 6.6 million sheep are raised, with production systems ranging from intensive dairy breeds in the South to extensive grazing systems in mountainous and marginal areas. The country recognises more than 50 sheep breeds, including both cosmopolitan and highly localised ones, which together represent a reservoir of biodiversity. Among them, industrial dairy breeds such as Lacaune are internationally renowned for Roquefort cheese production, while numerous local meat or hardy breeds—such as Basco-Béarnaise, Manech Tête Noire, or Noire du Velay—play a central role in maintaining regional production systems and landscapes.

The Ouessant breed is primarily raised on an island in Brittany, the Soay breed lives on the isolated Soay Island off the coast of Scotland, and the Black Belly breed originates from the West Indies. These breeds may exhibit specific genetic characteristics due to their island location.

Genomic resources for sheep have steadily improved over the past decade. The first reference genome assembly, Oar_v3.1, was published in 2014 (Jiang et al., 2014), followed by major updates. Assembly contiguity improved in ARS-UI_Ramb_v2.0 (Davenport et al., 2022), and the Y chromosome of a Churro sheep was added in ARS-UI_Ramb_v3.0 in 2023, which is the current reference assembly. A telomere-to-telomere assembly (T2T-sheep1.0) was released in 2024 (Luo et al., 2025).

SNP chips, such as the OvineSNP50 BeadChip (Kijas et al., 2009), have enabled genome-wide association studies (GWAS), genomic selection (Meuwissen et al., 2001), and large-scale diversity analyses, including the Sheep HapMap and subsequent international efforts (Kijas et al., 2012). More recently, short-read whole-genome sequencing projects, such as the International Sheep Genomics Consortium datasets, have provided millions of variants across hundreds of breeds, but they still depend on existing reference assemblies. The next announced release (run3) contains 3522 animals whose metadata are already available (Brauning, 2022).

As in other species, short-read sequencing has limitations, particularly for structural variant detection. Long-read technologies, which costs have recently dropped, now enable highly contiguous assemblies. The SeqOccin project aimed at generating assemblies for a set of species, among which *Ovis aries*.

Twelve sheep breeds were selected to represent the diversity of (mostly) French *Ovis aries*: Basco-Béarnaise, Black Belly, Blanche du Massif Central, Corse, Ile de France, Lacaune, Manech Tête Noire, Noire du Velay, Ouessant, Rambouillet, Soay, and Vendéen. Those include the major French dairy breeds: Lacaune, Basco-Béarnaise, Manech Tête Noire, Corse, some meat breeds: Blanche du Massif Central, Ile de France, Noire du Velay, Vendéen, a wool-reference breed (Rambouillet), a tropical-origin breed (Black Belly) and two isolated breeds (Ouessant, Soay). These 12 new assemblies will complement existing references, support pangenome construction, and foster comparative studies across sheep populations. They will provide an essential resource for the identification of structural variants and for linking genomic diversity with phenotypic diversity in French and international sheep populations.

## Material and methods

### Genome information

The domestic sheep genome follows the Bovidae family chromosomal organisation with 26 autosomes and 2 sex chromosomes (2n = 54). As in other Bovidae, autosomes are acrocentric, meaning that their centromeres are located terminally. Sheep genome size has been estimated at around 2.97 Gb using flow cytometry and Feulgen densitometry, with values in the literature ranging between 2.67 Gb and 3.31 Gb depending on the method used. Using k-mer based estimation with Jellyfish (Marçais & Kingsford, 2011) and GenomeScope2.0 (Ranallo-Benavidez et al., 2020) on PacBio HiFi data, genome size was estimated at ∼2.9 Gb with low heterozygosity (<0.3%) and ∼37% repeated sequences. The reference genome Oar_rambouillet_v2.0 (NCBI, GCA_016772045.1) has a total size of 2.86 Gb, 82.9 Mb contig N50, 141.1 Mb scaffold N50, and ∼96.2% complete BUSCO score. This assembly was used as a reference to scaffold our assemblies without Hi-C in chromosomes and to check all assembly metrics.

### Sample collection & technical validation

Blood samples were collected by Genphyse unit, as allowed by the government under the agreement number: 31113050. The Qiagen Tips 100 was used to extract high molecular weight DNA from theses fresh blood sheeps. DNA was stored at ™20°C in TE buffer (10 mM Tris–Cl, pH 7.5; 1 mM *EDTA*). For short read sequencing, a home made protocol was used. Genomic DNA concentrations were measured using the Qubit fluorimetry system (Life Technologies) with the High Sensitivity(HS) kit for detection of double-stranded DNA (Thermo Fisher, Part #Q32854). Fragment size distributions were assessed using the Femto pulse DNA 165kb Kit (Agilent).Purity was measured using a Nanodrop system (Thermofisher). All the samples were purified with AMPureXP beads (Beckman Coulter, A63882) to obtain correct ratios i.e. 260/280: 1.8–2 and 260/230:2-2.2.

### Sequencing

All sequencing was performed at the GeT-PlaGe core facility at INRAE Toulouse (GeT-PlaGe, 2022).

At each step, DNA was quantified using the Qubit HS kit. DNA purity was tested using a Nanodrop and size distribution and degradation assessed using the Femto pulse DNA 165kb Kit or Fragment Analyser kit. Purification steps were performed using AMPure PB beads (PacBio 100-265-900).

#### PacBio Sequel II

Library preparation was performed according to the manufacturer’s instructions “Procedure & Checklist Preparing HiFi SMRTbell Libraries using SMRTbell Express Template Prep Kit 2.0”(100-938-900).

For CLR library (simple assembly): 10µg of DNA was purified then sheared at 60kb using the Megaruptor3 system (Diagenode). Using SMRTbell Express Template prep kit 2.0, a Single strand overhangs removal, a DNA and END damage repair step were performed on 10µg of sample. Then blunt hairpin adapters were ligated to the library. A size selection step using a 30kb cutoff was performed on the BluePippin Size Selection system with “0.75% DF Marker U1 high pass 30–40kb VS3” protocol. A 70kb library was recovered. Using Binding kit 2.0 kit (101-789-500) and sequencing kit 2.0 (101-820-200) with 1 h of annealing and 4h of binding, the primer V4 annealed and polymerase 2.0 bounded library was sequenced by diffusion loading onto 1or2 SMRTcells 8M (101-389-001) on Sequel II instrument at 50 to 70 pM with a 15hours movie. A coverage of minimum 30X was obtained per sample.

For HiFi library (Lacaune assembly): 20µg of DNA was purified then sheared at 15 kb using the Megaruptor3 system (Diagenode, N°E07010003). Using SMRTbell Express Template prep kit 2.0, a Single strand overhangs removal, a DNA and END damage repair step were performed on 10 llµg of sample. Then blunt hairpin adapters were ligated to the library. The library was treated with an exonuclease cocktail to digest unligated DNA fragments. A size selection step using a 10 kb cutoff was performed on the BluePippin Size Selection system to obtain a 20 kb library. Using Binding kit 2.0 kit and sequencing kit 2.0 with 1h for annealing and 1h for binding, the primer V2 (101-847-900) annealed and polymerase 2.0 bounded library was sequenced by diffusion loading onto 4 SMRTcells on Sequel II instrument at 50 to 70 pM with a 2-hour pre-extension and a 30-hour movie. A coverage of 40X min was obtained.

#### Illumina paired-end WGS 2 × 150 bp

Parent whole genome reads used to phase the Lacaune assembly were sequenced following the protocol below.

DNA-seq libraries were prepared according to the Illumina protocol TruSeq Nano DNA High Throughput Library Prep Kit. DNA was fragmented by sonication on Pixul, size selection was performed using Sample Purification Beads of the library kit (ratio 1/1 beads water) and adaptators IDT for Illumina – TruSeq DNA UD Indexes were ligated before sequencing. Library quality was assessed using a Fragment Analyzer (Agilent) with High Sensitivity NGS Kit (DNF-474-0500). Sizes of 600 bp were obtained. Libraries were quantified by qPCR using the KAPA Library Quantification Kit (Roche, 07960140001). DNA was sequenced on 1/2 S4 NovaSeq 6000 lanes using a paired-end read length of 2 × 150 bp with the Illumina S4 Reagent kit (300 cycles). This sequencing produced a 70X coverage per sample.

#### Hi-C

Blood sample was processed using the Arima Hi-C Kit (ref. 510008) according to the manufacturer’s protocols. The library was prepared using the TruSeq DNA PCR-Free Kit and quantified by qPCR on QuantStudio 6 device (Applied Biosystems, ThermoFisher Scientific) using the Kapa Library Quantification Kit. Sequencing was performed in a 2 × 150 paired-end mode on a NovaSeq6000 (Illumina).

#### ONT

At each step, DNA was quantified using the Qubit HS kit. DNA purity was tested using the nanodrop and size distribution and degradation assessed using the Fragment analyzer (AATI) High Sensitivity DNA Fragment Analysis Kit. Purification steps were performed using AMPure XP beads (Beckman Coulter). Library preparation was performed according to the manufacturer’s instructions “1D gDNA selecting for long reads (SQK-LSK114)”. For 1 Flowcell, 5 µg of DNA was purified then sheared at 20 kb using the Megaruptor system (Diagenode). A 10 kb size selection step using the BluePippin Size Selection system was performed. A one-step DNA damage repair+ END-repair + dA tail (NEB) of double stranded DNA fragments was performed on 1 µg of sample. Then adapters were ligated to the library. The library was loaded on FLO-PRO004 (R10.4.1) flowcells and sequenced and PromethION instruments at 6/10 fmol within 72 h. Nuclease flush steps were done when necessary and possible, i.e. when only 30% of the pores were still in sequencing. We produced a 20 to 50X coverage.

### Genome assembly

As presented in the previous section, five types of reads were produced during the project. Three types of long reads were generated: PacBio CLR for Basco-Béarnaise, Corse, Ile De France, Black Belly, Blanche du Massif Central, Soay, Noir du Velay, Vendéen, Manech Tête Noire, Rambouillet and Ouessant breeds, PacBio HiFi for Lacaune and Oxford Nanopore for Soay and Black Belly. For the Lacaune animal only, Hi-C reads were used for scaffolding and parental Illumina short reads were used to phase haplotypes. The long read types were differing in error rate and therefore could not be assembled using a unique piece of software. Because of low throughput for Soay and Black Belly, ONT reads were produced two years later and a third assembly strategy was then set up using up to date software packages.

The first « High quality » strategy, used for the Lacaune breed only, was based on HiFi reads assembled with hifiasm version 0.16.1 (Cheng et al., 2021) and parental kmer dictionaries (-1 and -2 parameters) to separate haplotypes on the chromosome level. The kmer dictionaries were produced with yak count version 0.1 (Li, 2026) and a kmer size of 37. The haplotype separation was checked with merqury version 1.3 (Rhie et al., 2020) using default parameters (for the corresponding results section : Lacaune_contig_merqury.hapmers.blob.png image). Before running merqury the parental reads were used to produce a kmer dictionary with meryl version 1.3 (Miller et al., 2008) with default parameters.

The second “Medium quality” assembly strategy was based on PacBio CLR reads which were assembled for six breeds with flye version 2.9 (Kolmogorov et al., 2020) with the --pacbio-raw and --genome-size 3.0g parameters. Because of the low read quality this strategy includes a contig polishing round using the same long reads which was performed with ArrowGrid-release 489ebf8 from SMRTLink version 10.2 (Chin et al., 2013). Briefly the reads were realigned onto the contigs and the software used the sequencing signal of all the reads at all positions to decide which nucleotide to put into the consensus.

Because Soay and Black Belly DNA samples did not sequence well with PacBio CLR new DNA preparations were resequenced with Oxford Nanopore Technologies. In this third strategy CLR plus ONT reads were assembled by pecat ((Nie et al., 2024; lemene, 2026)) unzip version 0.0.3 using default parameters which first corrected the input reads by aligning them all versus all to generate corrected reads. These reads were then assembled and separated in two haplotypes which were polished independently. This last strategy produced two haplotypes per breed.

### Genome scaffolding

Two scaffolding strategies were used. The first one for Lacaune, based on Hi-C reads in which both haplotypes were scaffolded independently. First, the reads were aligned to each haplotyped contig set with juicer version 1.5.6 (Durand et al., 2016) with default parameters. The resulting Hi-C link file was processed with 3d-dna release 529ccf4 with parameter –s 0 in order to stop scaffolding at the first round. The resulting scaffolding was manually curated using juicebox version 1.11.8 to produce chromosomes. Chromosomes were also checked with merqury.

All the other breed assemblies were scaffolded on the Ramb_v2.0 reference genome from the NCBI website (https://ftp.ncbi.nlm.nih.gov/genomes/all/GCF/016/772/045/GCF_016772045.1_ARS-UI_Ramb_v2.0/GCF_016772045.1_ARS-UI_Ramb_v2.0_genomic.fna.gz) with ragtag version 2.0.1 using minimap2 version 2.24.(Li, 2018) Briefly, after aligning the contigs to the reference chromosomes, alignment uniqueness and quality enabled to decide contig location and orientation and contigs were then merged and separated with a spacing sequence of 1000 Ns. Non-matching or non-uniquely matching contigs were kept as isolated in the resulting fasta file.

### Assembly validation

Several metrics were produced to validate the scaffolded assemblies. For all of them assemblathon_stats.pl script was run to extract contig and scaffold statistics including total length, N50, L50 and GC content. Busco scores (Manni et al., 2021) were also produced in all cases using version 5.1.2 with mammalia_odb10 database. For high quality and corrected reads, kmer QV and completeness were calculated using merqury version 1.3 (Rhie et al., 2020) with default parameters and a blob graph was generated for the Lacaune breed, using parental reads kmer content.

## Results

The produced reads have been used to build a high-quality haplotypes assembly for the Lacaune breed and 13 other assemblies for the Basco-Béarnaise, Black Belly, Blanche du Massif Central, Corse, Ile de France, Manech Tête Noire, Noire du Velay, Ouessant, Rambouillet, Vendéen, and Soay breeds including two haplotypes for Black Belly and Soay.

The metrics below highlight the high quality of this data.

**Table 1:** Sheep data volume. Data volume for each breed. The first column corresponds to the species, the second to the number of reads, the third to the total read size, the fourth to the average read length, the fifth to the sequencing coverage, and the last one to the EBI/EMBL identifiers.

| Breed | Read numbers | Total size | Average length of reads | Coverage (X) | ID EBI/EMBL |
| --- | --- | --- | --- | --- | --- |
| Basco-Béarnaise | 12 390 118 | 210 755 663 146 | 17 010 | 78 | ERR14939018, ERR14939019 |
| Black Belly | 15 681 879 | 214 506 283 916 | 13 679 | 79 | ERR14939038, ERR14939040, ERR15094490 |
| Blanche du Massif Central | 8 797 142 | 244 778 603 184 | 27 825 | 90 | ERR14939031, ERR14939034 |
| Corse | 6 346 421 | 187 694 432 996 | 29 575 | 69 | ERR14939024 |
| Ile De France | 13 071 305 | 252 457 228 479 | 19 313 | 93 | ERR14939020, ERR14939021 |
| Lacaune | 8 955 257 | 155 092 908 126 | 17 319 | 57 | ERR14939025, ERR14939026, ERR14939027, ERR14939028, ERR14939030 |
| Manech Tête Noire | 6 268 742 | 129 697 397 847 | 20 689 | 48 | ERR14939023 |
| Noire Du Velay | 11 560 649 | 350 943 758 467 | 30 356 | 129 | ERR14939014, ERR14939017 |
| Ouessant | 15 572 499 | 234 868 374 416 | 15 082 | 86 | ERR14939015, ERR14939016, ERR14939029 |
| Rambouillet | 3 951 022 | 161 001 235 378 | 40 749 | 59 | ERR14939033 |
| Soay | 13 983 580 | 128 234 211 951 | 9 170 | 47 | ERR14939032, ERR14939035, ERR14939036 |
| Vendéen | 5 467 414 | 135 300 243 452 | 24 746 | 50 | ERR14939022 |

**Table 2:** Sheep Assembly statistics. Evaluation per breed of assembly quality and comparison with the reference (GCF_016772045.1_ARS-UI_Ramb_v2.0_genomic.fna.gz). Column 1 gives the breed (and haplotype if present), column 2 the corresponding EBI/EMBL ID, column 3, the number of scaffolds obtained using the assembler, column 4, the total assembly length, column 5 and 6, the scaffold N50 and L50. Columns 7,8,9 correspond to the metrics generated on contigs.

| Breed | ID EBI/EMBL | Number of scaffolds | Total size of scaffolds | N50 scaffold length | L50 scaffold count | Number of contigs | N50 contig length | L50 contig count |
| --- | --- | --- | --- | --- | --- | --- | --- | --- |
| Basco-Béarnaise | GCA_984514235.1 | 2441 | 2 651 381 193 | 101 116 994 | 8 | 3487 | 11 764 114 | 69 |
| Black Belly haplotype 1 | GCA_984506455.1 | 1035 | 2 722 815 387 | 95 689 921 | 9 | 1338 | 23 221 762 | 37 |
| Black Belly haplotype 2 | GCA_984506415.1 | 1819 | 2 712 954 963 | 94 820 913 | 9 | 2353 | 13 136 129 | 63 |
| Blanche du Massif Central | GCA_984507005.1 | 999 | 2 645 246 055 | 100 908 164 | 8 | 1499 | 27 076 389 | 30 |
| Corse | GCA_965363785.1 | 1533 | 2 655 464 741 | 95 071 006 | 8 | 2106 | 19 576 709 | 39 |
| Ile De France | GCA_984514435.1 | 2868 | 2 654 071 811 | 101 223 162 | 8 | 5561 | 29 803 54 | 266 |
| Lacaune haplotype 1 | GCA_984506435.1 | 384 | 2 778 774 854 | 94 408 239 | 9 | 421 | 77 725 785 | 12 |
| Lacaune haplotype 2 | GCA_984506445.1 | 326 | 2 898 238 363 | 97 125 495 | 9 | 393 | 84 416 955 | 12 |
| Manech Tête Noire | GCA_984506815.1 | 2320 | 2 663 399 558 | 101 168 904 | 8 | 3430 | 11 856 191 | 67 |
| Noire Du Velay | GCA_984514685.1 | 939 | 2 643 891 132 | 101 251 129 | 8 | 1384 | 31 079 500 | 27 |
| Ouessant | GCA_984506805.1 | 4678 | 2 674 701 900 | 101 528 116 | 8 | 6984 | 4 171 537 | 190 |
| Rambouillet | GCA_984513465.1 | 726 | 2 647 152 022 | 100 917 375 | 8 | 1060 | 36 955 596 | 20 |
| Soay haplotype 1 | GCA_984506635.1 | 960 | 2 684 727 422 | 101 330 441 | 8 | 1997 | 11 711 593 | 69 |
| Soay haplotype 2 | GCA_984506465.1 | 398 | 2 437 909 743 | 91 163 615 | 8 | 984 | 8 667 548 | 85 |
| Vendéen | GCA_984538235.1 | 1057 | 2 651 323 919 | 100 805 136 | 8 | 1687 | 22 502 704 | 37 |
| Reference Rambouillet | GCA_016772045.1 | 142 | 2 628 146 905 | 101 274 418 | 8 | 226 | 43 178 051 | 24 |

**Table 3:**
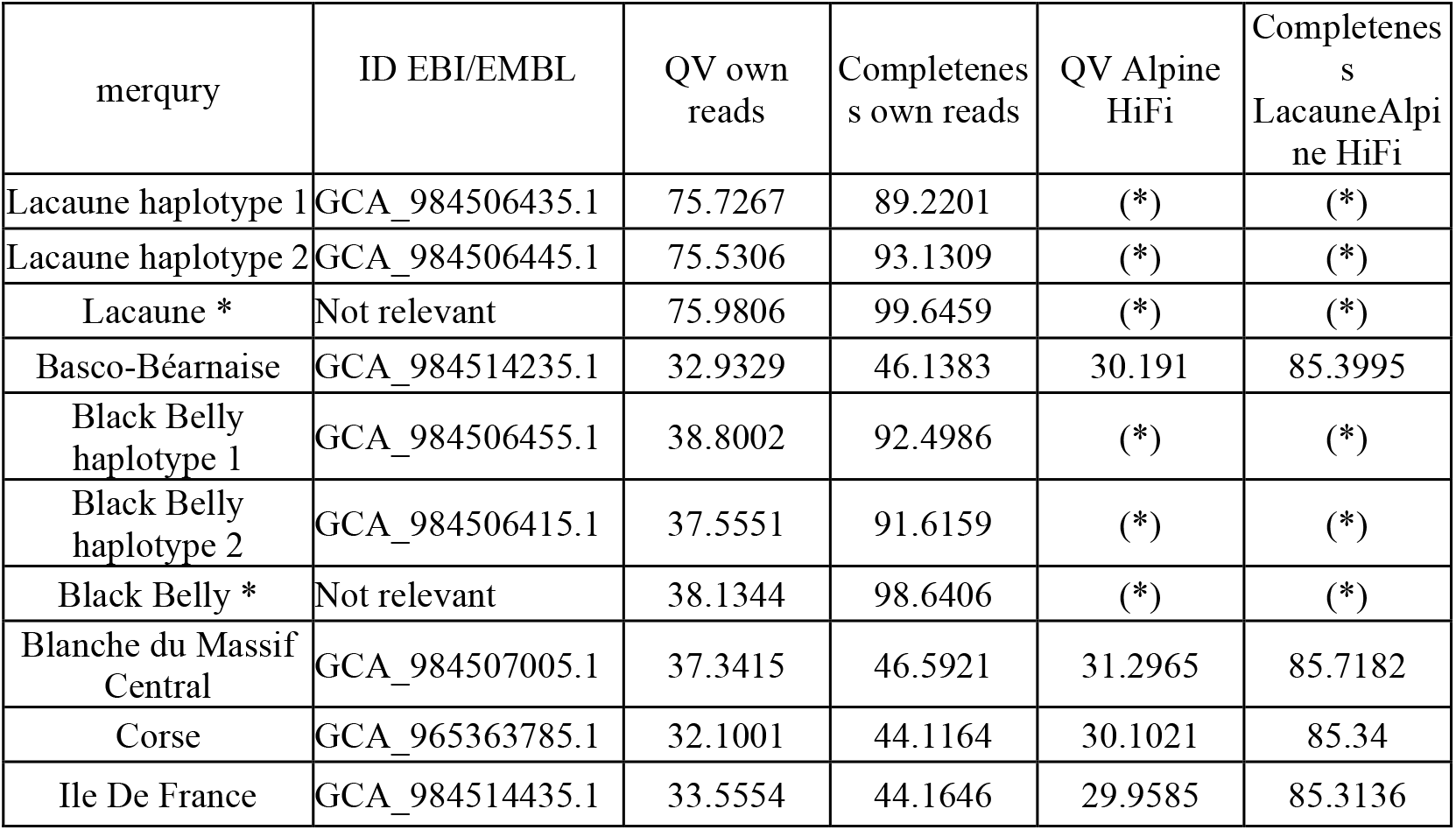
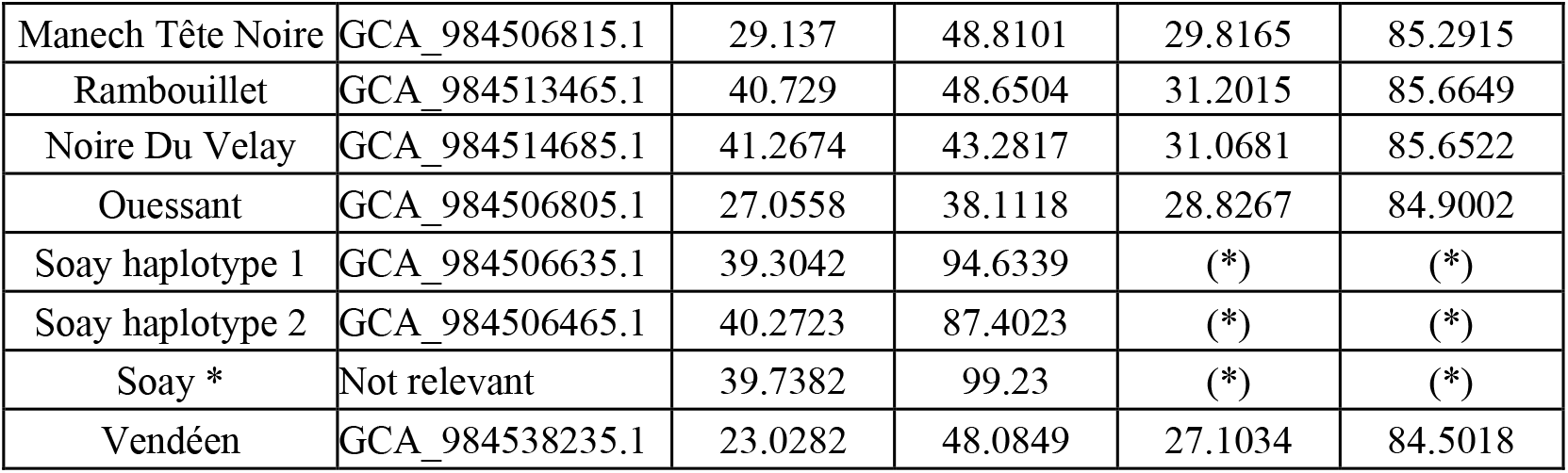
Merqury. (*)Not calculated for HiFi and ONT reads For each assembly the table presents merqury QV and completeness results. Column 1 gives the breed (and haplotype if present), column2 gives the assembly ID, column 3 and 4 present QV and completeness calculated using the reads with which the assembly was performed. Because CLR reads are of low quality (close to 12% errors) these metrics are poor. Therefore column 5 and 6 present the same metrics, but calculated with the Lacaune HiFi reads. (*) Both previous haplotypes were used.

**Figure 1.**
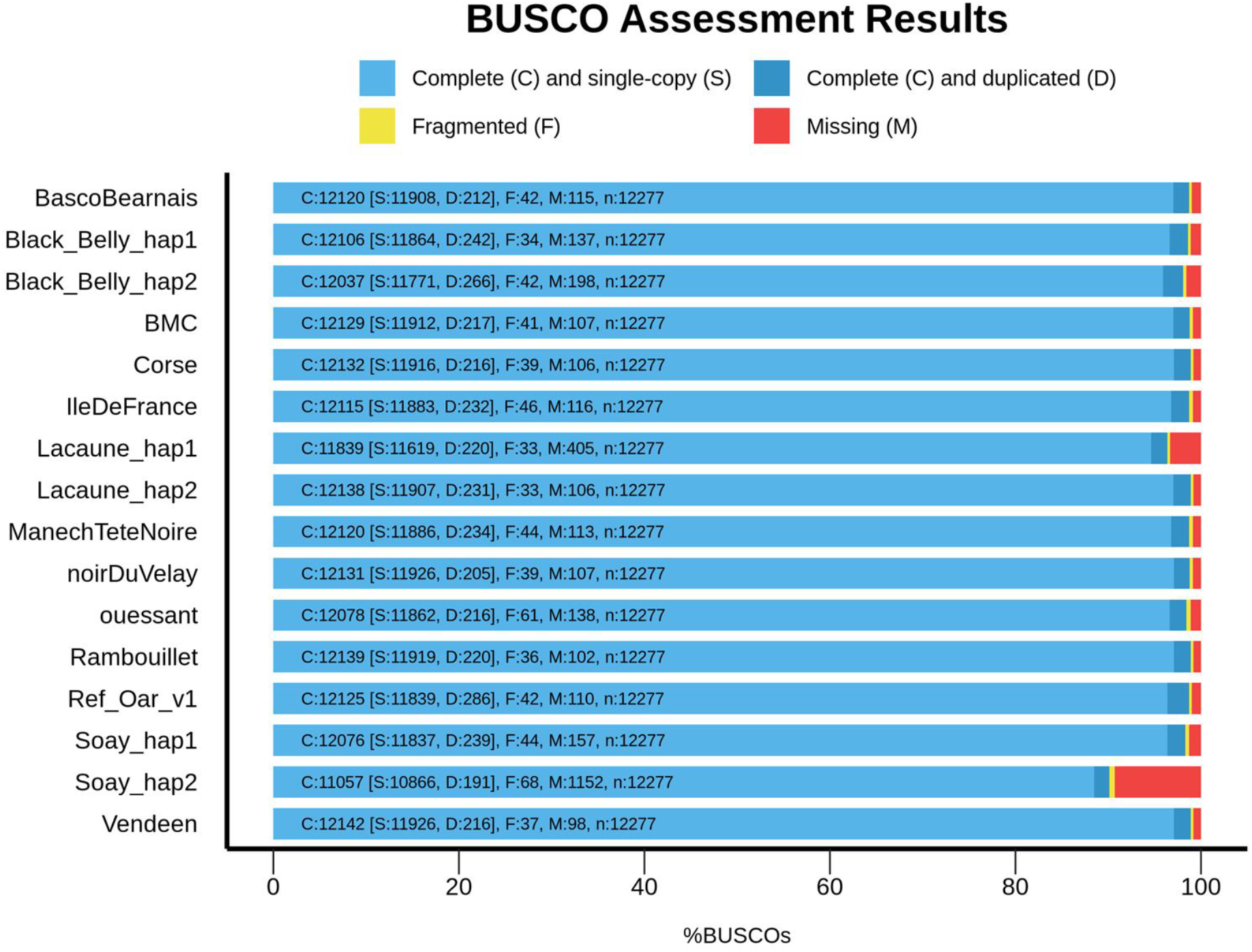
BUSCO. BUSCO plot of different *Ovis aries* assemblies. Summary plot produced with BUSCO V5.1 and mammalia_odb10 dataset.

## Acknowledgements

We thank all the people, who selected the breeds and collected the blood samples. The following list is probably not exhaustive:

- Stéphane Fabre, Jean-Michel Astruc and other researchers, engineers and technicians from the Unité Mixte de Technologie STAR (https://idele.fr/umt-star/).
- breeders, veterinarians and technicians of the relevant breeding organisations or INRAE experimental units for the following breeds: Basco-Béarnaise and Manech Tête Noire (CDEO, https://www.cdeo64.fr), Black Belly (*P3R*, INRAE, 2026, Pôle de Phénotypage Petits Ruminants https://doi.org/10.15454/1.5483259352597417E12), Blanche du Massif Central and Noire du Velay (ROM Selection, https://www.races-ovines-des-massifs.com), Corse (OS Brebis Corse, https://www.casgiucasanu.fr/category/races-locales), Ile de France (OSON, https://oson-gof.com, Lacaune (*UEF*, INRAE, 2026, Unité Expérimentale de la Fage https://doi.org/10.15454/1.548325523466425E12, Lycée agricole La Cazotte https://la-cazotte.educagri.fr, Service Elevage - Confédération Générale de Roquefort https://www.s-elevage.fr/Service_Elevage), Ouessant (GEMO, http://www.moutons-ouessant.com), Rambouillet (Bergerie nationale, https://www.bergerie-nationale.fr), Vendéen (OES Mouton Vendéen https://moutonvendeen.com) and Dr Susan Johnston, Institute of Evolutionary Biology University of Edinburgh, UK, for Soay sheep.

We thank “La Région Occitanie” and European Union for funding the project as part of the Occitanie Region’s “Regional Research and Innovation Platforms” call for projects under the FEDER-FSE MIDI-PYRENEES ET GARONNE 2014-2020 Operational Program.

GeT core facility (https://doi.org/10.15454/1.5572370921303193E12) is supported by France Génomique National infrastructure, funded as part of “Investissement d’avenir” program managed by the French Agence Nationale pour la Recherche (contract ANR-10-INBS-09). We are grateful to the genotoul bioinformatics platform Toulouse Occitanie (Bioinfo Genotoul, https://doi.org/10.15454/1.5572369328961167E12) for providing computing and storage resources.

## Data, scripts, code, and supplementary information availability

Data are available online:

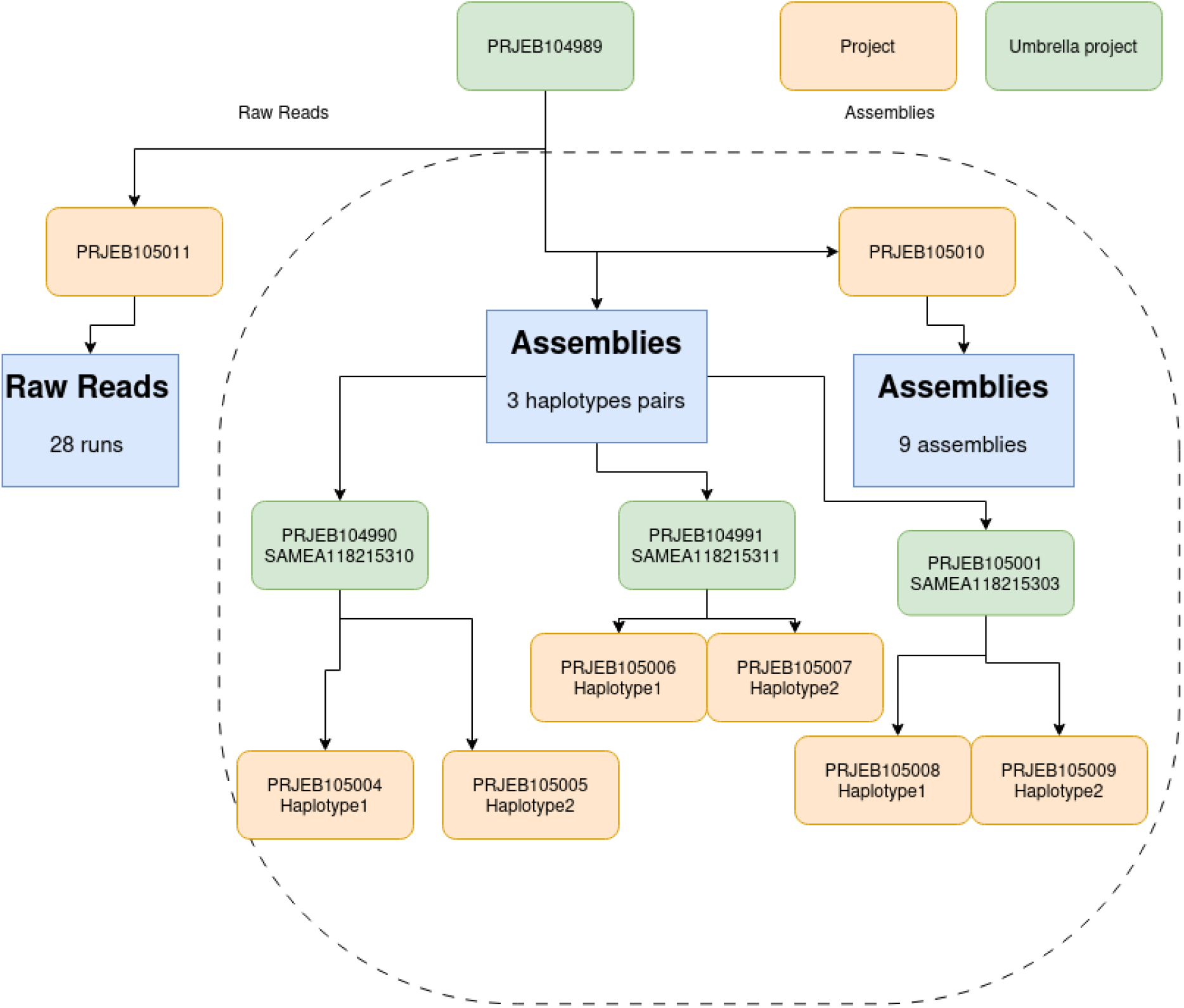

SWHID for the software packages used in this project. These IDs enable to download the software packages from https://archive.softwareheritage.org/

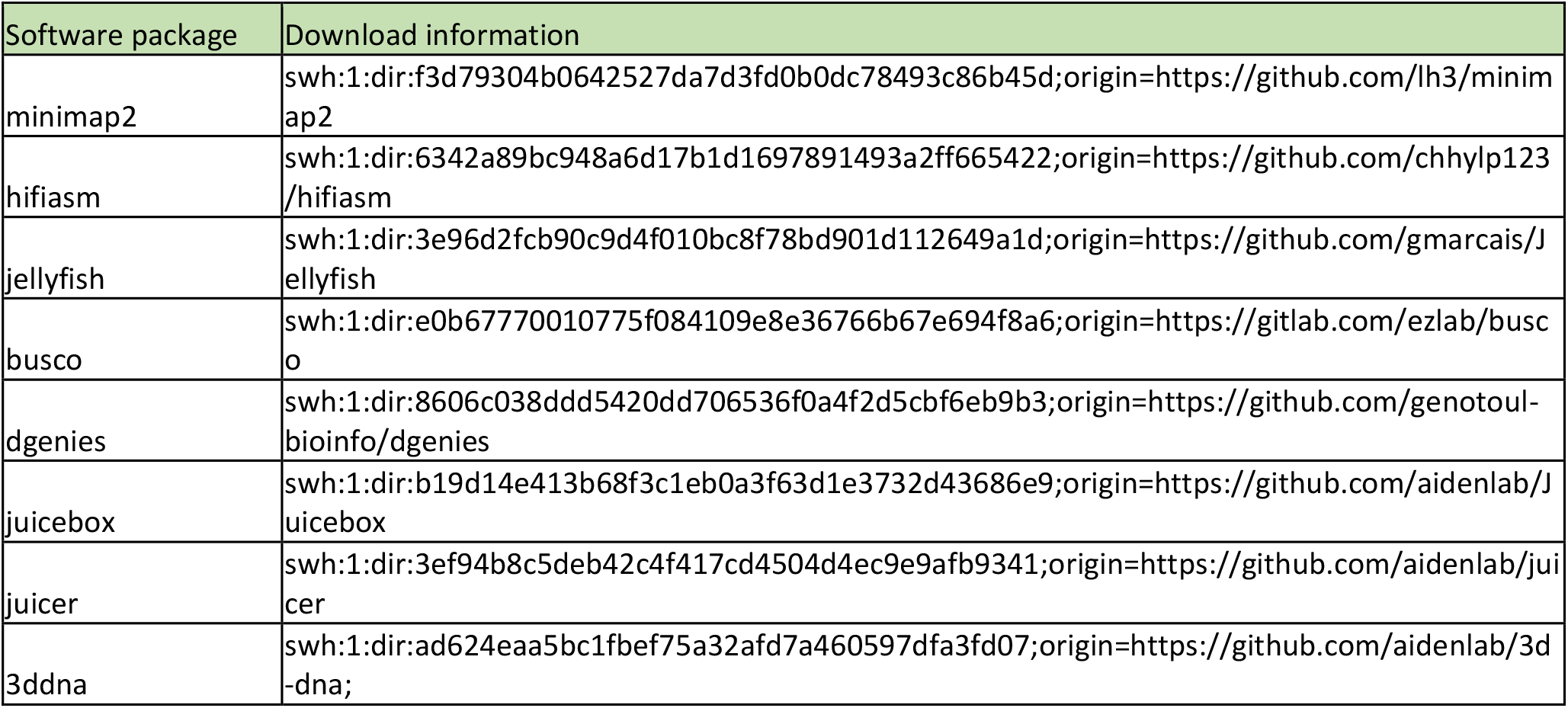
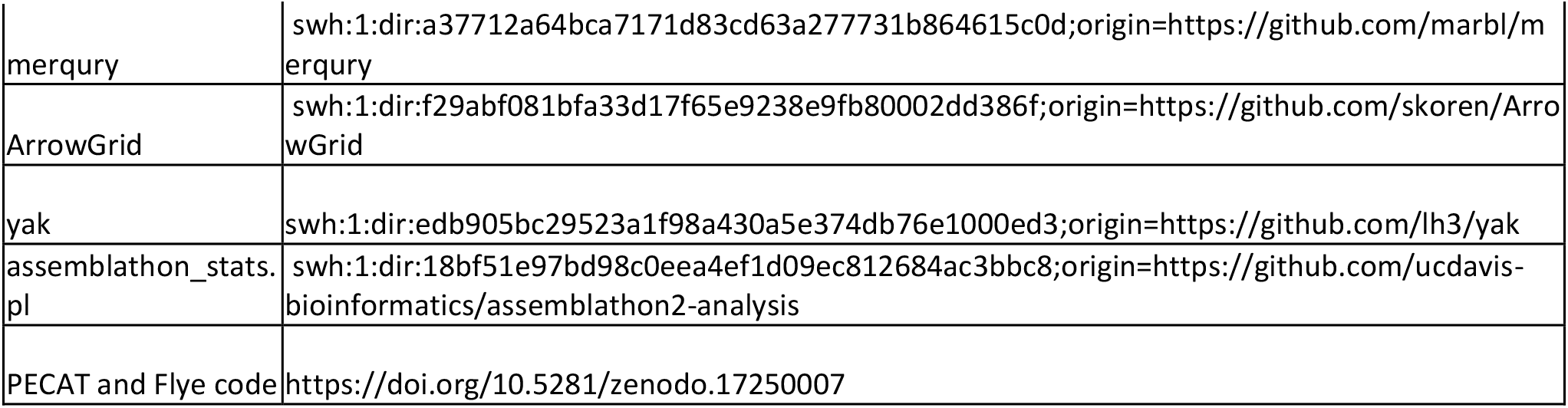

